# Galectin-8 is a major ligand of LILRB4 prompting MDSC functions in the tumor microenvironment

**DOI:** 10.1101/2022.07.27.501694

**Authors:** Yiting Wang, Yufan Sun, Shouyan Deng, Teng Song, Yungang Wang, Jie Xu

## Abstract

The LILRB4 myeloid receptor has been implicated in an immunosuppressive microenvironment, with specific antibodies under preclinical or clinical development for tumor immunotherapy. However, it remains largely unknown which natural ligand may trigger LILRB4 to expand myeloid derived suppressive cells (MDSC), and the relevant downstream signaling pathways are also under debate. Here we show that Galectin-8 is a high-affinity functional ligand of LILRB4, and its ligation induces MDSC by activating STAT3 as well as inhibiting NF-κB. Importantly, Galectin-8 but not APOE could induce MDSC, and both ligands bind LILRB4 in a non-competitive manner. Antibodies recognizing a defined epitope on LILRB4 could efficiently block Galectin-8 binding and neutralize its effects on MDSC induction and relevant signaling pathways. Galectin-8 expression promoted B16 tumor growth in mice, and knockout of LILRB4 attenuated tumor growth in this context. The LILRB4-specific Galectin-8 blocking antibody efficiently suppressed MDSC expansion and tumor growth *in vivo*. These results identify Galectin-8 as a functionally important ligand of LILRB4, highlighting the blockade of LILRB4-Galectin-8 interaction as a promising strategy for cancer immunotherapy.

## Introduction

Although T cell checkpoint immune therapies have brought huge success in the past decades, only about 30% of patients with specific cancer type showed significant and durable response to treatment [1, 2]. The complexity of tumor microenvironment adds up to a suppressive phenotype of immune cells, and preceding clinical research have shown that high level of tumor infiltrating myeloid cells correlates with poor outcome of ICB [3, 4]. Mechanistic investigations identified a group of myeloid derived cells that are immune suppressive and able to limit the potency of T cell ICB therapies [5]. Thus, targeting myeloid derived suppressive cells (MDSC) suggests a compelling approach to deal with the inefficiency of T cell ICBs [6, 7].

MDSC represents a heterogeneous population, making it hard to identify an amendable target. RNA-sequencing of the two subgroups of MDSC, monocytic MDSC (M-MDSC) and granulocytic MDSC (G-MDSC), identified CD33 as a common marker on MDSCs, regardless of tumor type. According intervention approaches targeting CD33 have been found to restore T cell proliferation and CAR-T activity [6]. However, the expression and essential functions of CD33 in bone marrow and other normal tissue may limit the translational potential of CD33-targeting strategy.

Recently, attention has been drawn to the leukocyte immunoglobulin-like receptors (LILRBs), a family of inhibitory receptors identified in 1997 [8]. Their roles in MDSC expansion and functions have been increasingly evident and emphasized [9]. LILRB4 had long been regarded as an orphan receptor, until APOE was identified as its functional ligand in 2018 [10]. APOE was reported to bind LILRB4 on the surface of acute myeloid leukemia (AML) cells, promoting cancer progression and inhibiting T cells. Therapeutical antibody was developed accordingly [11]. Beyond AML, LILRB4 was evidently related to poor prognosis in solid tumor [12]. Researchers discovered that LILRB4 plays an important role in promoting MDSC expansion and cancer progression [13-15], making LILRB4 a promising target for a variety of tumor types [16, 17]. However, it is still unclear which ligand of LILRB4 may be involved in shaping the microenvironment of solid tumors, which is irrelevant to the functions of previously reported ligands or binding proteins such as APOE, CD166 and β-amyloid[13]. Typically, one receptor may bind different ligands, each inducing distinct functional effects [18] [19, 20]. Thus, a functionally relevant ligand of LILRB4 involved in maintaining an immunotolerant microenvironment has been in hot pursuit in the field.

Galectins are a family of revolutionarily conserved proteins that bind glycans, with potential roles in cancer cell survival, angiogenesis, metastasis and immune modulation [21-24]. Galectin-9, a ligand of TIM-3, was found to induce expansion of mMDSC and resistance to PD-1 blockade in lung cancer patients [25-27]. Both Galectin-9 and Galectin-8 (Gal-8) were identified as prognostic factors in cervical cancer and many other cancer types[28-30]. Gal-8 has been found to be upregulated in prostate cancer [31], promoting cancer cell migration by binding CD166 [32, 33]. In the present study, we report Gal-8 as a high-affinity, agnostic ligand of LILRB4, and characterize its roles in expanding MDSCs and maintaining a tolerant tumor microenvironment. We also demonstrate the targeting the Gal-8/LILRB4 immune checkpoint signaling by blocking antibodies can suppress tumor growth.

## Results

### Galectin-8 is related to myeloid cell mediated suppression in tumor microenvironment

Galectin-8 firstly attracted our attention by its association with T cell dysfunction in the tumor microenvironment, as characterized by the Tumor Immune Dysfunction and Exclusion (TIDE) model used for predicting response to immune checkpoint blockade [34]. As shown in **Fig.1A**, in tumors expressing low level of Gal-8, T cell infiltration associated with better prognosis. However, such an association was absent in tumors highly expressing Gal-8 (**Fig.1A**). Among the immunosuppressive cell types that may associate with T cell exclusion, MDSC was ranked on top (**Fig.1B**). To further explore Gal8-8’s role in MDSC, we used TIMER to calculate the fraction of MDSC infiltration and performed Pearson correlation analysis between MDSC infiltration and LGALS8 expression based on TCGA datasets [35]. The correlation analysis between LGALS8 expression and MDSC infiltration was positive in a variety of cancer types, including ACC, SKCM (**Fig.1C**), ESCA, KIRP, LIHC and UCS (**Supplementary Fig.1B**). Consistently, Gal-8 showed no direct effect on CD4+CD25+FOXP3+ Tregs (**Supplementary Fig.1A**). Consistantly, we further performed phagocytosis assay on THP-1-derived macrophages treated with or without Gal-8 for 24 hours (**Fig.1D**), which confirmed a role of Gal-8 in decreasing the phagocytosis of macrophages (**Fig.1E**).

**Figure1.**
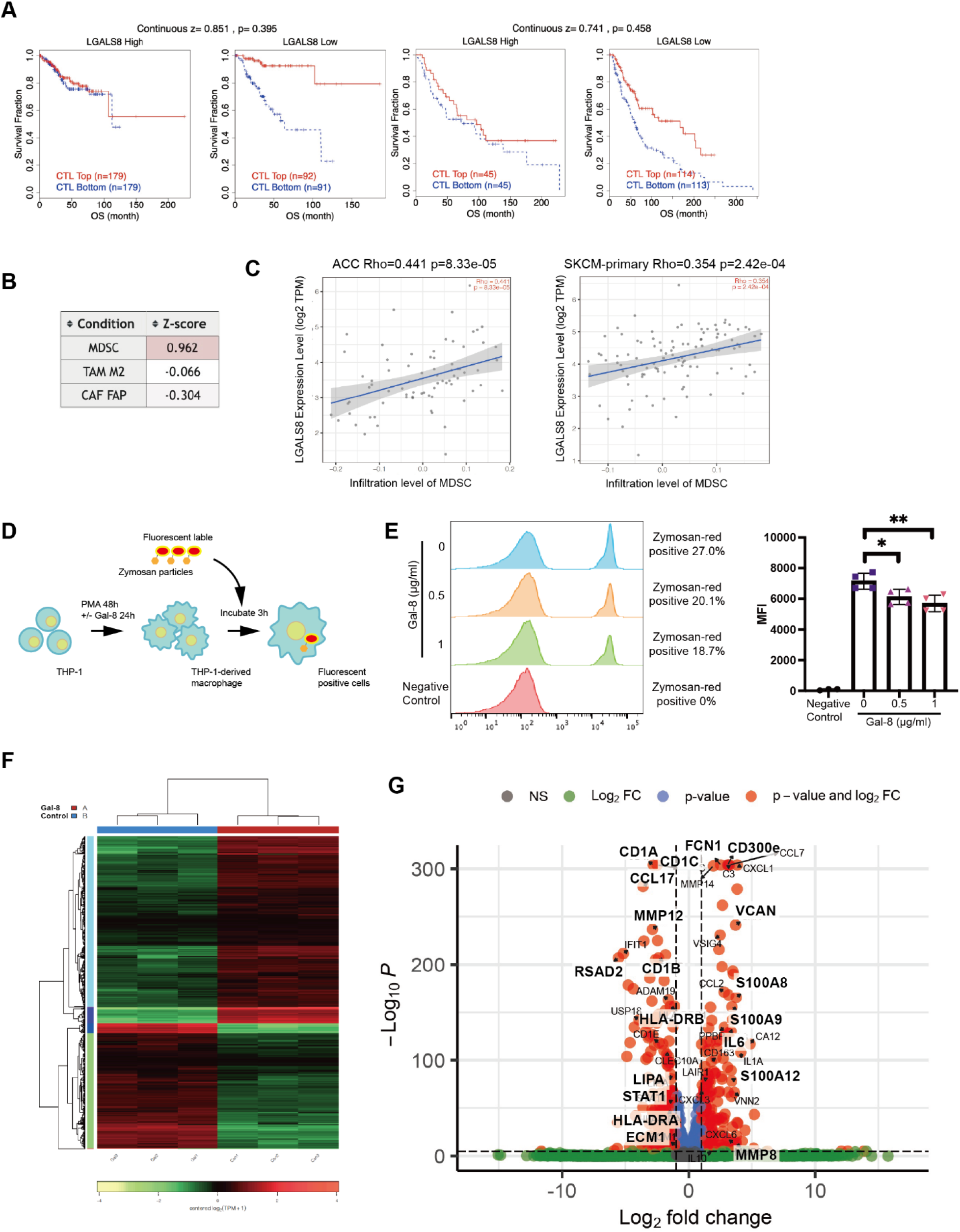
Galectin-8 is related to myeloid cell mediated suppression in tumor microenvironment. (A) Data analysis with TIDE shows that LGALS8 plays an important role in T cell dysfunction in tumor microenvironment. (B) Spearman correlations between Galectin-8 expression and TILs. (C) TIMER was used to calculated MDSC fraction and correlation with LGALS8expression in the indicated types of tumors from the TCGA dataset. (D) Schematic procedure of phagocytosis assay. (E) Phagocytosis assay (FC) shows that Galectin-8 inhibits the ability of phagocytosis. F. Transcriptome analysis of CD14+ monocytes exposed with Galectin-8. (F) Heatmap of transcriptome sequencing data. Samples of 2 groups including control and gal-8 were analyzed. (G) Volcano plot of transcriptome sequencing data. Significantly regulated genes were labeled.

Human CD14+ monocytes sorted from PBMC were grouped and treated with or without Gal-8, followed by transcriptome sequencing analysis. The results demonstrated clear difference in gene expression pattern between two groups (**Fig.1F**). Among the most significantly upregulated genes, C300e, IL6 and MMP8 have been reported to be critical for MDSC functions, and S100A8/9/12, FCN1 and VCAN represent markers of MDSC phenotypes (**Fig.1G**). Many other genes associated with MDSC functions and induction were upregulated, including CXCL1/2/5, CCL2/7, C3, MMP14, FPR1, IL1A, MERTK, APQ9, IL10 et al (**Supplementary Table.1**). Meanwhile, HLA-DR, a negative marker of MDSC, as well as genes negatively related to MDSC accumulation and expansion, such as MMP12, RSAD2, LIPA, STAT1 and ECM1, were significantly downregulated (**Fig.1G**). Some other downregulated genes were related to DC and macrophage immune response and T cell activation, such as CD1a/b/c, CCL17 et al (**Supplementary Table.2**).

These observations consistently suggested that Gal-8 may be involved in the suppressive phenotype of monocytes. According to by the Single Cell Expression Atlas [36], Gal-8 is highly expressed in the testis (spermatids) and placenta (trophoblasts), both being immune privileged organs (**Supplementary Fig.1B**).

### Galectin-8 binds LILRB4 to induce MDSC expansion

In support to the correlation between Gal-8 and MDSCs, strong interaction was also found between Gal-8 and LILRB4 (**Fig.2A**), a receptor driving MDSC functions [12, 15, 37, 38]. Immunofluorescent labeling of Gal-8 overexpressed in cells displayed cytoplasmic punctate-like distribution, as shown in **Fig.2B**. However, in cells co-expressing LILRB4, the distribution of Gal-8 was drastically changed to a similar pattern as that of LILRB4 (**Fig.2B** and **Supplementary Fig.2A**). The co-immunoprecipitation assay also confirmed interaction between both proteins in cells (**Fig.2C**). We then characterized the binding affinity between Gal-8 and LILRB4 by ELISA and Biolayer Interferometry (BLI) assay. When Gal-8 was coated at 2 μg/ml on ELISA plate, the EC50 of LILRB4 binding to Gal-8 was 1.38 μg/ml with R square of 0.994 (**Fig.2D**). The Biolayer Interference (BLI) assay determined the binding affinity between Gal-8 and LILRB4 (KD=1.02μM) (**Fig.2E**), being several folds higher than that of PD-1/PD-L1 (varied between 7.2 to 8.2 μM depending on assays) [39]. Crosslinking of LILRB4 and Galectin8 showed abundant formation of binding complex (**Supplementary Fig.2B**).

**Figure 2.**
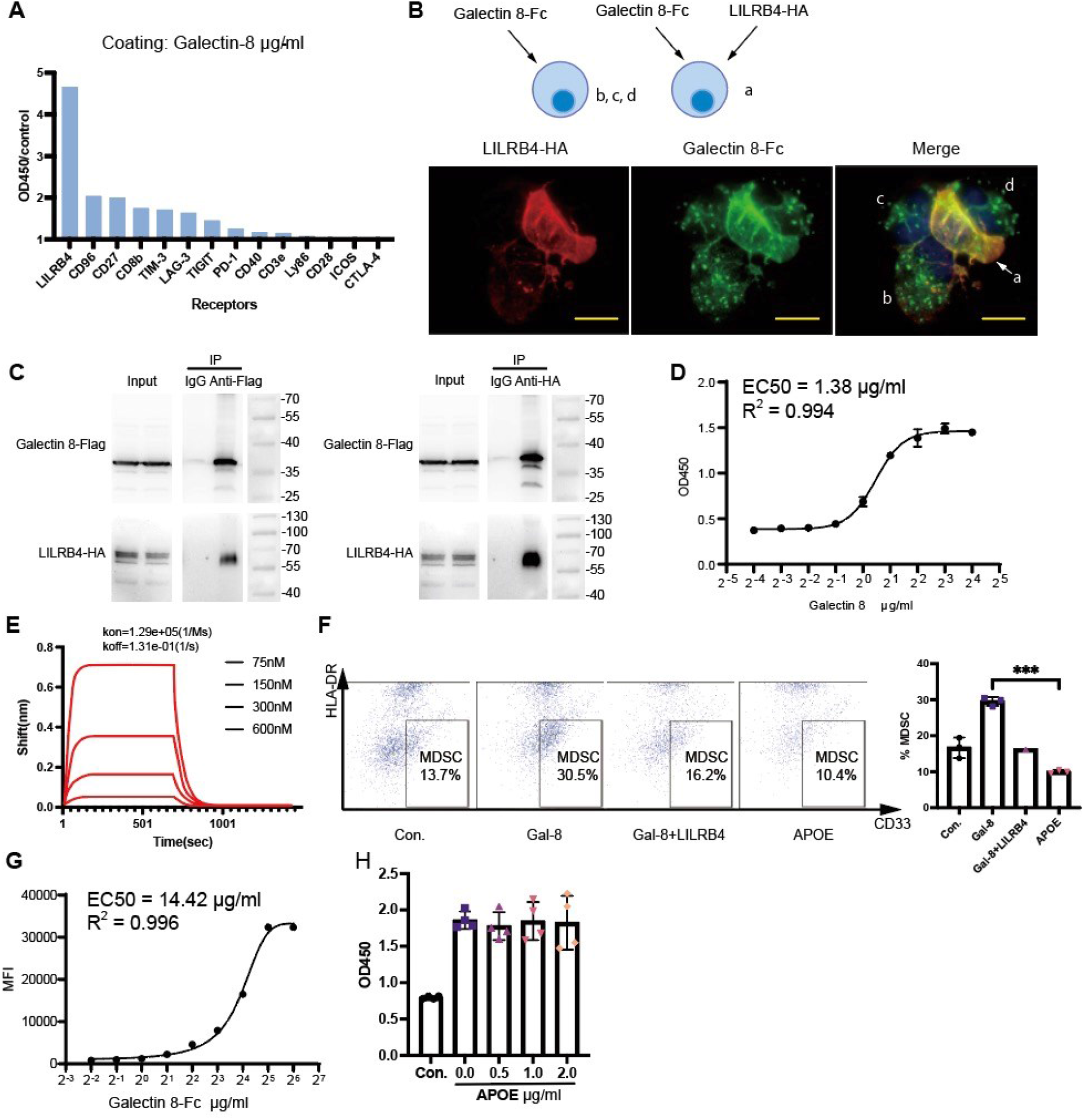
Galectin-8 binds LILRB4 to induce MDSC expansion. (A) ELISA screening of potential Galectin-8 interactors revealed LILRB4. (B) 100X microscopic views of Immunofluorescence showing Galectin-8 binds to cell surface LILRB4. (C) CoIP of Galectin-8 and LILRB4. (D) Analysis of EC50 from ELISA test. Galectin-8 was coated by concentration scale, LILRB4-Fc was added to 1 μg/ml. (E) BLI assay showing the association-disassociation curve between Galectin-8 and LILRB4 with the kinetics constants. (F) Flow cytometry showing the fractions of MDSCs in PBMCs incubated with Gal-8, Gal-8 plus LILRB4 ectodomain, and APOE. (G) Flow cytometry showing the binding of Galectin8 to LILRB4 expressed on cell surface. (H) ELISA showing the binding between Galectin-8 and LILRB4 in presence of different concentrations of APOE.

Consistent with potential myeloid regulating ability of Gal-8, LILRB4 was reported to induce tolerance of dendritic cells and expansion of MDSCs [13, 15]. Thus, we investigated the effect of Gal-8 on to MDSCs defined as CD11b+, CD33+ and HLA-DR-cells. As expected, Gal-8 expanded the MDSC population, while the presence of soluble LILRB4 counteracted the effect (**Fig.2F**). In contrast to Gal-8, APOE displayed no effect on MDSCs (**Fig.2F**).

The different effects on MDSCs of Gal-8 and APOE suggest that both ligands may bind LILRB4 by different conformational regions or manners. To test this, we first confirmed the binding of Gal-8 to LILRB4 expressed on HEK293 cell surface (**Fig.2G**), resembling the reported ability of APOE [10]. When an increment concentration of APOE was added, the binding of Gal-8 to LILRB4 was not affected (**Fig.2H**), confirming different sites involved in their binding.

### Gal-8-LILRB4 interaction activates STAT3 and inhibits NF-κB through SHP-1

We then went to characterize the downstream signaling of Gal-8/LILRB4 axis involved in MDSC regulation. The intracellular domain of LILRB4 contains three ITIMs that may recruit Src homology 2 (SH2)-containing tyrosine phosphatases (SHPs) and SH2 domain containing inositol phosphatase (SHIP) to transduce inhibitory signals [40]. There is also evidence showing potential positive or negative regulation of monocyte functions by LILRB4, depending on the positions of tyrosine residues in its ITIMs [41]. For a better demonstration of Gal-8/LILRB4 function, a LILRB4-knockdown (KD) THP-1 cell line was constructed using shRNA lentivirus (THP-1 shLILRB4), and cargo lentivirus was used to constructed a control cell line (THP-1 NC). Different concentrations of Gal-8 was added to THP-1 (control or LILRB4-KD) cells, followed by Western Blot detection of phosphorylation of downstream phosphatases. Interestingly, SHP-1 (but not SHP-2 or SHIP-1) was significantly phosphorylated in control THP-1 cells treated by Gal-8. However, in LILRB4-KD THP-1 cells, the phosphorylation of SHP-1 was substantially attenuated (**Fig.3A**). This finding agreed with the previous report on the ability of LILRB4 in promoting SHP-1 phosphorylation [42].

**Figure 3.**
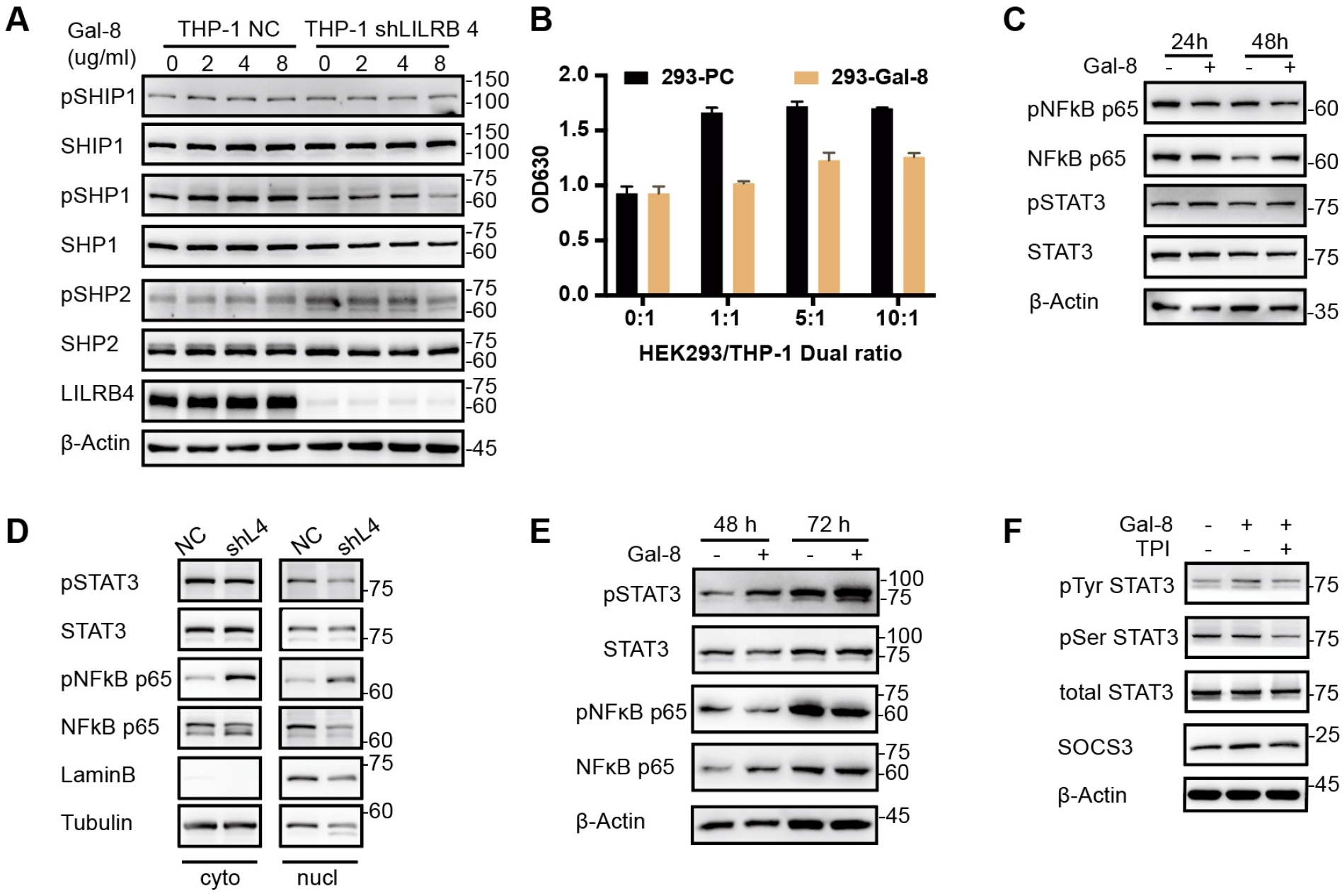
Gal-8-LILRB4 interaction activates STAT3 and inhibits NF-κB through SHP-1. (A) Phosphorylation of SHP1 was upregulated by the interaction of Galectin8 and LILRB4 in THP-1 cells. (B) THP-1 NFκB reporter cells cocultured with Galectin-8-overexpressed 293 cells rather than control 293 cells showed lower activation of NFκB signal. (C) Immune blotting demonstrates Gal-8 inhibited NFκB while activated STAT3 in THP-1 cells. (D) Nuclear protein extraction of NC/shLILRB4 THP-1 cells and immune blotting revealing effect of LILRB4 on NFκB and STAT3. (E) Immune blotting of human CD14+ cells. (F) Immune blotting of SHP-1 inhibitor, TPI, inhibited STAT3 phosphorylation in human CD14+ cells.

Although it has been reported that APOE-LILRB4 interaction activated NF-κB through SHP-2 in Acute myeloid leukemia cells [10], we found that Gal-8 played a distinct role. By coculturing Gal-8 overexpressing HEK293 cells with THP-1 (NF-κB) reporter cells, we found that Gal-8 significantly inhibited NF-κB signaling (**Fig.3B**).

It has been found that SHP-1, SHP-2 and SHIP may adapt LILRB4 ITIMs to different downstream signaling pathways. Indeed, the Gal-8/LILRB4/SHP1 axis in myeloid cells displayed no effect on ERK1/2 signaling (**Supplementary Fig.3A**), although LILRB4 on lung cancer cells may activate ERK1/2 through SHP-2 and SHIP1 [43]. Likewise, the Gal-8/LILRB4/SHP1 axis did not affect Akt phosphorylation (**Supplementary Fig.3A**), which may be a result if LILRB4 recruits SHP-2 in certain lymphoma cells [44].

However, we could detect substantial NF-κB inhibition upon Gal-8/LILRB4 interaction (**Fig.3C**), possibly due to its specific effect on SHP-1. In addition, STAT3 was reported to be activated by SHP-1 [45], so was it involved in MDSC induction [46]. Accordingly, STAT3 phosphorylation was tested and proven activated by Gal-8 (**Fig.3C**). To ensure that Gal-8 took effect through LILRB4, we used LILRB4 KD THP-1 cell as comparison. As a result, NF-κB phosphorylation level was higher and could not be inhibited by Gal-8 and induced STAT3 phosphorylation was weaker in LILRB4 KD cells (**Supplementary Fig.3B**). Specifically, tyrosine phosphorylation was induced rather than serine phosphorylation in STAT3.

Questions may be raised here, as Deng and colleagues discovered decreased NF-κB nuclear translocation in LILRB4 KO THP-1 cell [10]. This intriguing question compelled us to repeat the test, in order to functionally confirm the activity of these two transcription factors. We detected NF-κB and STAT3 after separation of cytoplasm and nuclear fractions. Surprisingly, the result was consistent with what was reported that NF-κB in nuclear protein pool decreased in LILRB4 KD cells, but our findings was also confirmed that phosphorylated NF-κB was vastly enhanced without LILRB4 (**Fig.3D**). Meanwhile, phosphorylation of STAT3 was lower in LILRB4 KD cells than control cells (**Fig.3D**).

The regulation of STAT3 was further investigated in sorted CD14+ cells (Fig.3E). Phosphorylation of STAT3 elevated with Gal-8 treatment. In addition, a selective inhibitor of SHP-1 (TPI) successfully blocked STAT3 phosphorylation (**Fig.3F**). This finding supported the function of SHP1-STAT3 pathway in monocytes.

### Gal-8-LILRB4-SHP-1 inhibits ADAM17 through TRAF6-NFκB

Afterwards, we explored the mechanism of NF-κB inhibition. By dephosphorylating TRAF6, SHP-1 was reported to inhibit TRAF6 ubiquitination [47], and TRAF6 ubiquitination leads to activation of NF-κB [48, 49]. Therefore, TRAF6 was pulled down with antibody and protein G beads and its ubiquitination was detected. In LILRB4 KD cells, ubiquitiniation of TRAF6 was stronger and phosphorylation of NF-κB was promoted (**Fig.4A**). In wildtype THP-1 cells, Gal-8 treatment down regulated ubiquitination of TRAF6 (**Fig.4B**).

**Figure 4.**
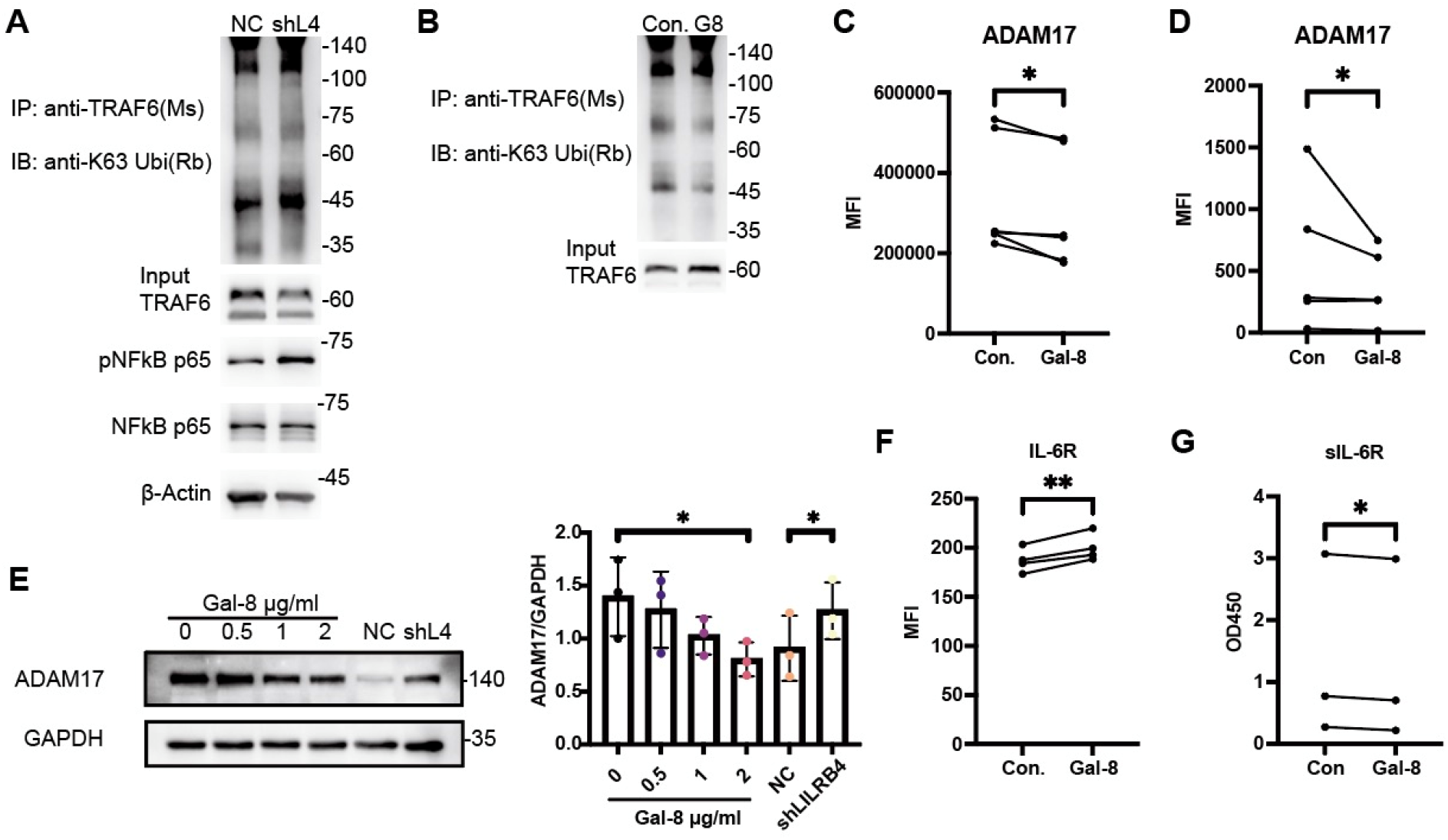
Gal-8-LILRB4-SHP-1 inhibits ADAM17 through TRAF6-NFκB. (A) IP of THP-1 NC/shLILRB4 cells by anti-TRAF6 antibody detected with anti-K63 Ubi antibody. (B) IP of Gal-8 treated/untreated THP-1 cells by anti-TRAF6 antibody detected with anti-K63 Ubi antibody. (C) Flow cytometry showing that Gal-8 downregulated ADAM17 in THP-1 cells. (D) Flow cytometry showing that Gal-8 downregulated ADAM17 in human monocytes. (E) Immune blotting showing ADAM17 expression alteration in THP-1 cells and statistical analysis. (F) Flow cytometry showing that Gal-8 upregulated ADAM17 in THP-1. (G) Soluble IL-6R decreased by Gal-8 tested by ELISA.

As a well-addressed yet complicated transcript factor, NF-κB controls transcription of a variety of target genes. Through literature exploration, we found a NF-κB regulating factor, ADAM17[50], that is closely related to IL-6 signaling [51]. ADAM17 was believed to participate in immune regulation [52]. Membrane expression of ADAM17 on THP-1 cells was revealed to be downregulated by Gal-8 (**Fig.4C**), the same with human PBMC sorted monocytes (**Fig.4D**). In consistence with flow cytometry assay, whole cell expression level of ADAM17 decreased with increased Gal-8 (**Fig.4E**). On the contrary, knockdown of LILRB4 increased ADAM17 expression. ADAM17 was reported to cleavage membrane proteins into soluble forms, thus cutting down on their membrane expression [51]. In our experiments, PD-L1 [53] and CD163 [54] was not notably affected (**supplementary Fig.4A&B**). Of note, membrane expression of IL-6R elevated with decrease of ADAM17 (**Fig.4F**). Meanwhile, soluble IL-6R decreased in culture medium (**Fig.4G**). As expected, upregulation of membrane IL-6R empowers IL-6 signal transduction and reduced sIL-6R strengthens the effect. Consequently, the activation of STAT3 and MDSC expansion is fueled.

### Galectin-8 and LILRB4 interaction promotes tumor growth in vivo

Before carrying out in vivo experiment, we confirmed binding of mouse LILRB4 and human Galectin 8 by ELISA (**Fig.5A**). We applied CRISPR/Cas9 technique on C57BL/6J mice to build lilrb4 knockout (HE) mice. Wildtype C57BL/6J mice was used as control group. B16, a melanoma cell line derived from C56BL/6 mice, was transfected with plasmid to constructed stable Galectin 8 overexpressing cell line. Plasmid control was used for control cell line construction. The same procedure was performed on MC38, a C56BL/6 derived colon cancer cell line. Overexpression (OE) and wildtype (WT) cells were injected subcutaneously to lilrb4 knockout (KO) and/or wildtype (WT) mice (**Fig.5B**).

**Figure 5.**
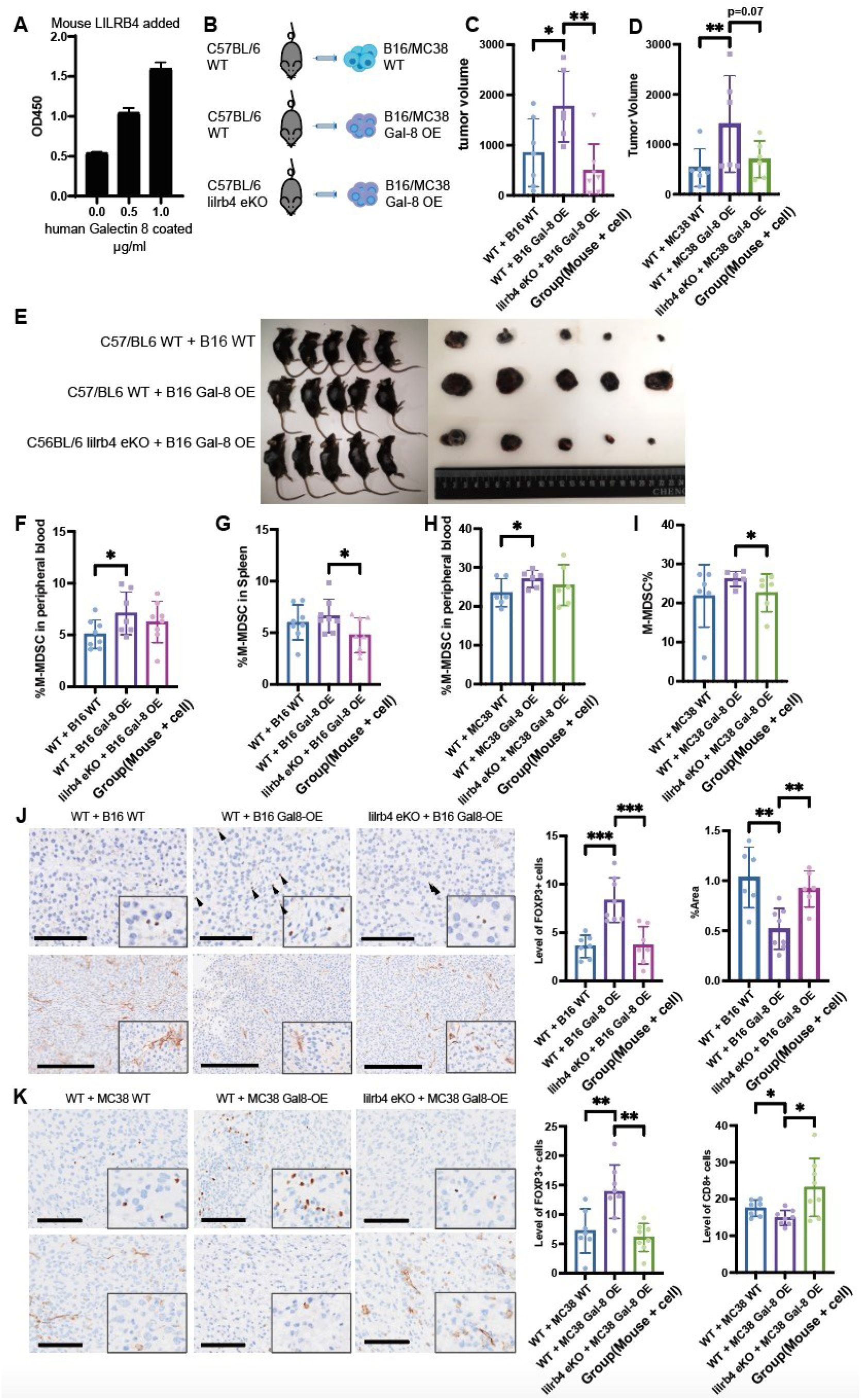
Galectin-8 and LILRB4 interaction promotes tumor growth in vivo. (A) ELISA result of plate bound human Gal-8 and soluble mouse LILRB4. (B) Strategic diagram of tumor transplant mice model. (C) B16 tumor volume 15 days after transplantation of 2e5 cells per individual. (D) MC38 tumor volume 28 days after transplantation of 1.5e6 cells per individual. (E) Photo of B16 tumor in vivo and ex vivo. (F) Ratio of MDSC in peripheral blood of mice bearing B16 tumor. (G) Ratio of MDSC in spleen of mice bearing B16 tumor. (H) Ratio of MDSC in peripheral blood of mice bearing MC38 tumor. (I) Ratio of MDSC in spleen of mice bearing MC38 tumor. (J) FOXP3+ cells stained in B16 tumor and statistical analysis (Bar=100 um). CD8+ cells stained in B16 tumor and statistical analysis (Bar=250 um). (K) FOXP3+ cells stained in MC38 tumor and statistical analysis (Bar=100 um). CD8+ cells stained in MC38 tumor and statistical analysis (Bar=100 um).

Specifically, 3e5 B16 cells OR 1.5e6 MC38 cells were injected per individual on day 0. Subcutaneous tumor was detectable in most of the mice from day 5 for B16 and from day 7 for MC38. Since then, the size of tumor was measured three times a week, until the tumor size was not ethically acceptable. As time went, Gal-8 OE tumor grew faster than WT tumor and faster in WT mice than in KO mice (**Supplementary Fig.5A&B**). Gal-8 OE tumor in WT mice outgrowing that in KO mice strengthened that Gal-8 induced immune suppression and tumor progression through binding with LILRB4 (**Fig.5C&D**).

After tumor-bearing mice were sacrificed, dissected tumor were recorded photographically in vivo and ex vivo (**Fig.5E** and **Supplementary Fig.5C**). Periphera blood mononuclear cells (PBMCs) and solenocytes were sampled for Flow cytometry in order to gate and measure the level of M-MDSC cells, defined as CD11b+, Ly-6C+, Ly-6G-cells. In both splenocytes and PBMCs, the WT + Gal-8 OE group presented highest level of M-MDSC both in B16-bearng mice (**Fig.5F&G**) and MC38-bearing mice (**Fig.5H&I**). Notably, M-MDSC level of WT + Gal-8 OE was greater than that of WT + WT group in PBMCs (**Fig.5F**). While in splenocytes, M-MDSC level of WT + Gal-8 OE group outnumbered that of lilrb4 KO + Gal-8 OE group (**Fig.5G**). MC38 bearing mouse models share similar pattern of M-MDSC level (**Fig.5H&I**). It is possible that M-MDSC in central immune organs is closely related to LILRB4 phenotype, while M-MDSC in peripheral blood is more susceptible to tumor Gal-8 expression, suggesting mutual influence of Gal-8 and LILRB4 on overall immune activity of tumor-bearing mice. In order to further demonstrate tumor infiltrating immune status, IHC staining of CD8 and FoxP3 was performed to tumor samples. FoxP3 positive T regulator cell (Treg) and MDSC are major components of suppresive tumor microenvironment. CD8+ T cells are representative of effector T cells. MDSC was known to suppress T cells and induce Treg [55]. Results supported that Gal-8 and LILRB4 upregulated Treg level and downregulated CD8+ T cell infiltration both in B16 tumor (**Fig.5J**) and in MC38 tumors (**Fig.5K**).

In recent years, LILRB4 was firstly regarded as a promising target for AML drug development. Shortly after, targeting LILRB4 in solid tumors began to draw enormous attention. To date, at least three LILRB4 antibody drugs have been uncovered to be going through preclinical or clinical studies. Last year, A group of researchers from Merck & Co., Inc. reported that LILRB4 antibody abrogated myeloid immunosuppression and enabled tumor killing [17], though the process and strategy of antibody screening was not revealed. Animal generated antibodies potentially bind numerous sites that may lead to different function. Without knowing a functioning ligand, it would be arduous to screen a potent antibody in early development stage.

Our findings identified a tumor related functional ligand for LILRB4, Gal-8. Gal-8 corelates with poor prognosis in patients with optimal CTL tumor infiltration, indicating that Gal-8 may induce immune suppression in tumor microenvironment. The interaction between Gal-8 and LILRB4 was confirmed by a variety of methods, so was their effect in inducing suppressive phenotype of monocytes. In vivo study emphasized their role in mediating immune suppression and tumor progression.

Mechanistic study unveiled two downstream pathways of LILRB4 in regulating monocyte activities. STAT3 was reported to be phosphorylated in activation of immature monocytes, which was thought to be a crucial step towards MDSC expansion in the two-stage model of proposed by D. I. Gabrilovich and colleagues [56]. In this model, activation of immature myeloid cells by STAT3 preceeds accumulation of suppressive myeloid cells, comprising two indispensible stages mutually contributing to MDSC expansion. Based on collective evidence, we believe that the activation of STAT3 through LILRB4 proves Gal-8 an unknown yet important myeloid suppressor.

On the other hand, decreased NF-κB phosphorylation was induced through LILRB4-SHP1-TRAF6 pathway. Even though NF-κB was reported by many to mediate immune suppression of MDSCs [57], its status during the induction of MDSC remained unclear. In macrophages, NF-κB was regarded as an activator of phagocytosis [58]. Accordingly, inhibition of phagocytosis was observed upon Gal-8 treatment. Therefore, NF-κB inhibition may be a regulator in monocyte suppression, which is one of the stages of MDSC development, as mentioned above. Interestingly, we managed to reveal ADAM17-IL6R pathways as a link between NF-κB inhibition and STAT3 activation. Collaboration of STAT3 and NF-κB pathways was assumed to drive cancer through cotroling communication betweencancer cells and inflammatory cells [59]. The interaction between two pathways adds more complexicity to the process of MDSC expansion. Actually, researchers have brought up assumptions supporting a multi-stage model of MDSC expansion [60].

In summary, STAT3 acts as an activator in immature myeloid cells, NF-κB plays a role as an inhibitory effector in cell behaviors such as phagocytosis. These two pathways induced by LILRB4 mutually enhance MDSC, showing that LILRB4 signaling is involved in multi stages of MDSC expansion. With its potential clinical value, the binding of Gal-8 and LILRB4 deserves more efforts and attention in drug developments.

## Methods and Materials

### In vivo Tumor models

All animal experiments were performed in strict accordance with the relevant ethical guidelines, approved by the department of laboratory animal science of Fudan University and the Institutional Animal Care and Use Committee of Renji Hospital, School of Medicine, Shanghai Jiaotong University. After one week of adaptation to the environment, each individual mouse (female/male, 6 weeks old) was randomized into four groups (n=8 for each group) and injected 1.5e6^6^ mouse colorectal MC38 cells or 3e5 mouse melenoma B16 stable cells subcutaneously in the right flank (the establishment of stable clones as described later). Since tumors became visible, the tumor sizes were recorded every 2-3 days using a vernier caliper and calculated with the formula 1/2 × A × a^2^ (A and a, respectively, denote the length and the width of the tumor). In accordance with the ethical guidelines, mice would be sacrificed once the tumor volume reached 2cm^3^ or ulcers happened. Samples were collected as described.

### Generation of lilrb4 KO mouse

C57BL/6J mice embryos were collected and CRISPER/Cas9 technique was applied to knockout lilrb4 from the embryos. The KO status was validated by PCR. The KO embryos were then cultured and fed to breed their next generations. First-generation mosaic mice were crossed with C57BL/6J wild-type mice to obtain heterozygous mice (HE), and heterozygous mice were crossed to obtain homozygous mice (HO). We continued to paired and bred these mice until enough HO individuals were obtained.

### Cell Culture

Human blood cancer THP-1 cells were purchased from ATCC. mouse colorectal cancer MC38 cells and melanoma B16 cells were purchased from Kerafast. luciferase-reporter THP-1 dual cells were purchased from Invivogen. PBMCs were purchased from Mt-bio, HEK293 cells were given from Zhigang Lu’s Lab (IBS, Fudan, Shanghai, China) and all mycoplasma free. THP-1 and B16 cells were incubated in RPMI-1640 (Meilunbio) with 10% FBS (Gibco). THP-1 cells were treated with PMA for 3days to generate THP-1 derived macrophages. Phagocytosis assay was carried out per manufacturer instructions (ab234054, Abcam). HEK293 and MC38 cells were incubated in DMEM (Meilunbio) with 10% FBS (Gibco). THP1-dual cells were incubated in RPMI-1640 (Meilunbio) with 10% FBS (Gibco) and supplemental selective antibiotics (Invivogen). All cells were cultured at 37°C under 5% CO2. PBMCs were cultured in AIM-V (Gibco). Human CD14+ cells were positively selected per manufacturer instructions (17858, Stem cell).

### Antibodies and Reagents

The primary antibodies for GAPDH (KC-5G5, KANGCHEN), Actin (KANGCHEN), Tubulin (Affinity), LaminB(CST), LILRB4 (GTX87582, GeneTex), Flag-tag (4793, CST), HA-tag (3724, CST), p-Akt (4060, CST), Akt (4691, CST), p-STAT3 (9145, CST), STAT3 (9139, CST), p-NF-κB p65 (3033, CST), NF-κB p65 (8242, CST), p-ERK (sc-81492, Santa Cruz Biotechnology), ERK (4695, CST), S100A8 (15792-1-AP, ProteinTech), S100A9 (26992-1-AP, ProteinTech), SOCS3 (ab16030, Abcam), TRAF6 (8028, CST), Polyubiquitin (5621, CST), PD-L1 (13684, CST), CD163(333602, biolegend) and ADAM17 (ab2051, Abcam) were all commercially available.

Reagents involving recombinant protein Galectin-8 (10301-HNAE; SinoBiological), APOE (APE-H5246; Acrobiosystems), CD3ε (10977-H02H; SinoBiological), CTLA-4 (CT4-H5255; Acrobiosystems), CD28 (CD8-H525a; Acrobiosystems), CD96 (TAE-H5252; Acrobiosystems), LAG-3 (LA3-H5255; Acrobiosystems), TIM-3 (TM3-H5258; Acrobiosystems), CD40 (CD0-5253; Acrobiosystems), ICOS (ICS-H5258; Acrobiosystems), OX40 (OX0-H5255; Acrobiosystems), TIGHT (TIT-H5254; Acrobiosystems), LY86 (10242-H02H; SinoBiological), LILRB4 (16742-H02H; SinoBiological), CD27 (CD7-H5254; Acrobiosystems), PD-1 (10377-H02H; SinoBiological), CD8b (11031-HCCH; SinoBiological) were also purchased from the indicated suppliers.

### Luciferase assay

The THP-1 dual cells were resuspended in test medium RPMI-1640 (Meilunbio) +10%FBS (Gibco) to 1×10^6^/ml. We then added 180 μl of cell suspension per well of a flat-bottom 96-well plate (Costar) with indicated concentrations of Gal-8 protein. For the groups that required activation, Pam3CSK4 (Invivogen) for was added simultaneously. We incubated the plate at 37°C under 5% CO2 for 18 hours. We then added 20 μl sample per well into a 96-well plate (Costar), then 180 μl Quanti-blue solution (Invivogen) into each well. The plate was placed in SpectraMax i3x and proceed measurement immediately. The whole process was protected from light.

### Protein-protein binding ELISA

The protein was diluted in coating buffer (Solarbio) to indicated concentrations and added 100 μl per well into a 96-well ELISA plate (Costar). We placed the plate at 4°C overnight. We washed the plate with PBST (Applied Cells, Inc.) five times and added 100 μl 5% BSA (VWR Life Sciences) into each well and incubated at 37°C for 90 minutes. We repeated washing and added 100 μl protein into each well and incubated at 37°C for 60 minutes. We repeated washing and added 100 μl associated secondary antibodies conjugated with HRP into each well and incubated at 37°C for 30 minutes. After 5 times PBST washing, we added 100 μl TMB buffer (Invitrogen) into each well and incubated at 37°C for 15 minutes, and added 50 μl stop buffer (Abcam) to terminate the reaction. The plate was placed in SpectraMax i3x and read at 450 nm.

### Flow cytometry

The cells (2×10^6^/ml) were added 100 μl per well a flat-bottom 24-well plate (Costar) with indicated concentrations of protein and incubated as described. After incubation, samples were washed with flow cytometry staining buffer (Invitrogen) three times. Then diluted fluorescent antibodies were added at suggested concentrations and incubated per manufacturer instructions. Next, after being washed with staining buffer three times, samples were analyzed them by MACSQuant16 (Miltenyi). FlowJo V10 was applied to analyze the data. Antibodies used in FACS including anti-CD11b APC (17-0118-42, Invitrogen), anti-CD33 PE (303404, Biolegend), anti-HLA-DR APC-Cy7 (307618, biolegend), anti-Ly-6C FITC (128006, Biolegend), anti-Ly-6G PE-Cy5.5 (127616, biolegend), anti-rabbit PE (406421, Biolegend) and anti-mouse AF488 (A21202, Invitrogen) are all commercially available.

### Plasmid construction and Establishment of stable cells

The expression vectors encoding pcDNA3.1-HA-LILRB4 and pcDNA3.1-Flag-Galectin-8 were generated by inserting synthesized complementary DNAs into the pcDNA3.1 vector. All plasmids were sequenced to confirm whether the designed mutation was present, without any other unwanted mutation. Ectopic Flag-tagged Galectin 8 or HA-tagged LILRB4 plasmid were transfected into the Mc38/B16/HEK294 cells with Fugene (Promega). A blank vector control was applied. After approximately two-week incubation supplemented with 200/350/600 μg/ml G418 (Gibco BRL) (based on cell types) with refreshing the medium every 2-3 days, the single colonies were picked and verified by immunoblots. An optimal clone was chosen and expanded. The lentiviruses for LILRB4 shRNA were purchased from GeneChem. The transfection was carried out per manufacturer instructions. After 96 hours, the medium was refreshed with RPMI-1640 complete medium containing 2 μg/ml puromycin (Invivogen). The medium was refreshed every 2-3 days for two weeks and identified the transfection efficiency by immunoblot.

### Immunoblotting and Nuclear and Cytoplasmic Extraction

For immunoblotting, cells were lysed with RIPA buffer (Beyotime) supplemented with 1% proteinase and phosphatase inhibitors cocktail (ThermoFisher Scientific). The collected cell lysates were centrifuged for 15min at 12000rpm (4°C). The supernatant was reserved and the protein concentration was determined by BCA Protein Assay Kit (ThermoFisher Scientific). 5×SDS-PAGE loading buffer (Applied Cells, Inc.) was diluted to 1× with protein sample and heating at 100°C for 8 min. The protein extracts were subjected to appropriate concentrations of SDS–PAGE for electrophoresis and transferred to PVDF membranes (Bio-Rad). Membranes were blocked with 5% bovine serum albumin (ThermoFisher Scientific) for one hour at room temperature, and then incubated with the primary antibodies overnight at four degrees. Membranes were incubated with secondary HRP-conjugated antibodies (KANGCHEN) at room temperature for one hour. Before and after the incubation, the membranes were washed five times with TBST and then examined by ChemiDoc imaging system (Bio-Rad). Extraction of nuclear and cytoplasmic protein was performed per manufacturer instructions (78833, Thermo) before immune blotting.

### Co-immunoprecipitation

cells were harvested and lysed with IP Lysis Buffer (ThermoFisher Scientific) supplemented with 1% cocktail of proteinase and phosphatase inhibitor and PMSF (ThermoFisher Scientific). The cell lysates were centrifuged for 2min at 12000rpm at four degrees. After DNase (QIAGEN) treatment for 16 min at room temperature, 16μL was removed from each sample (300μL) and mixed with 4μL 5×SDS-PAGE loading buffer (Beyotime) to serve as the input control, and the rest was incubated with 1μg primary antibody or IgG overnight at 4 °C with slow-speed rotating. Protein G Agarose beads (20398, ThermoFisher Scientific) were also incubated with 4% BSA for blockade overnight at 4 °C with slow-speed rotating. Each sample added with equal volume of beads were again slow-speed rotated at RT for 1h. The samples were then washed 3-4 times with PBS with high-speed rotation, and mixed with 30μL SDS sample buffer and heating at 100°C for 8 min. The samples were detected by western blot as described above.

### Immunofluorescence

The culture medium of cells seeded in 8-well chamber slides (C7182, Sigma) was removed, and the cells were washed twice with PBS and then fixed with 4% formaldehyde (28908, ThermoFisher) for 20 min. After being washed twice with PBS, cells were permeabilized and blocked with 0.2% Triton X-100 and 1% BSA in PBS simultaneously at room temperature for 1h. Then cells were incubated with primary antibodies at 4°C overnight. After PBS washing for five times, cells were incubated with secondary antibodies for 20 minutes at room temperature. Then after PBS washing for five times, the slide was sealed with DAPI and observed with a fluorescence microscope. The quantification of fluorescence intensity and the colocalization was analyzed by ImageJ software (version 2.0.0-rc-69/1.52p).

### Immunohistochemistry

Tissues samples were deparaffinized and rehydrated, and the antigen retrieval was carried out in citrate antigen retrieval solution. After blocking endogenous peroxidase with 3% H2O2 for 15 min and blocking with goat serum for 1 hour, tissue samples were incubated with primary antibodies overnight at 4°C, followed by incubation with biotin-conjugated secondary antibody at RT for 1 hour. DAB was used as chromogen and nuclei were counterstained with hematoxylin. CD8 staining was assessed using Image J and FOXP3 staining were manually counted in 5 random scope per slide.

### Kinetics assay

Fc-tagged LILRB4 ECD protein and protein A probes were applied in kinetics assay. His-tagged LILRB4 and His-tag probes were applied in epitope binning. Gator biolayer interferometry system and data processing platform were utilized per manufacturer’s instrctions (SNGC00070, ProbeLife, Lnc.).

### Antibody generation and Fab production

The process of animal immunization and mono-clone selection are as described. The fab production was carried out using Pierce Fa preparation Kit (Thermo) per manufacturer’s instructions.

### Statistical analysis

Column bar graphs and scatter plots were plotted using GraphPad Prism V.9. Values were mean±SD from three independent experiments. Two-sided Student’s t-test was applied for the comparison of two independent samples and one-way analysis of variance with post hoc test (Tukey) was applied to compare more than two groups. ImageJ (V.2.0.0-rc-69/1.52p) was applied for the data quantification of immunofluorescence and western blot analysis. The average fluorescence intensity was means±SD from three independent experiments. The co-localization factor (Pearson’s R value) were calculated using ImageJ (V.2.0.0-rc-69/1.52p) with the plugin ‘coloc2’ to evaluate the co-localization between two proteins. The number of samples assigned to each treatment was selected to provide sufficient statistical power to discern significant differences in different groups, based on prior experience with the experiment. The only data points excluded were those that were clear outliers due to technical problems in assays done in triplicate experiments. In this work, p value <0.05 was considered to be statistically significant.

## Data availability

The authors declare that all data supporting the findings of this study are available within the paper and its Supplementary Information files.

## Author contributions

YW, YS, SD, TS and YW performed experiments; YW and JX wrote the paper; JX conceived and supervised the study.

## Acknowledgements

This work was supported by National Natural Science Foundation of China (No: 82030104, 81874050, 81572326), Basic Research Projects of Shanghai Science and Technology Innovation Action Plan (20JC1410700); National Key R & D Program of China (2016YFC0906002, 2016YFC0906002), Tang Scholar (XJ), and Startup Research Funding of Fudan University.

## Notes

### Competing Interest Statement

The authors have declared no competing interest.

