## Supplementary Information for "Galectin-8 is a major ligand of LILRB4 prompting MDSC functions in the tumor microenvironment"

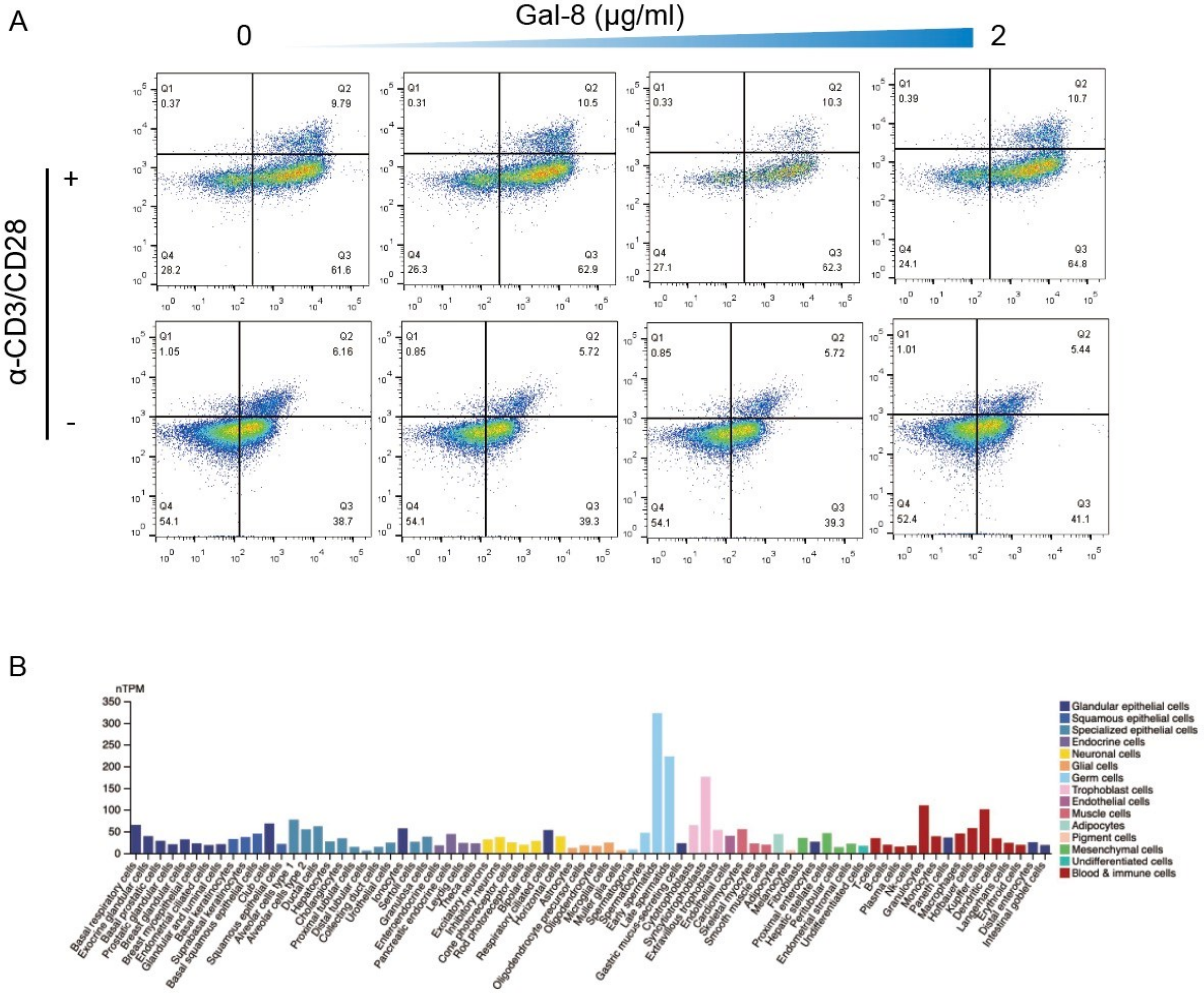


**Supplementary Figure 1.** (A) Gal-8 had no significant influence on Portion of CD4+CD25+FoxP3+ Treg. (B) Protein Atlas databases revealing Gal-8 enrichment in different single cell types.

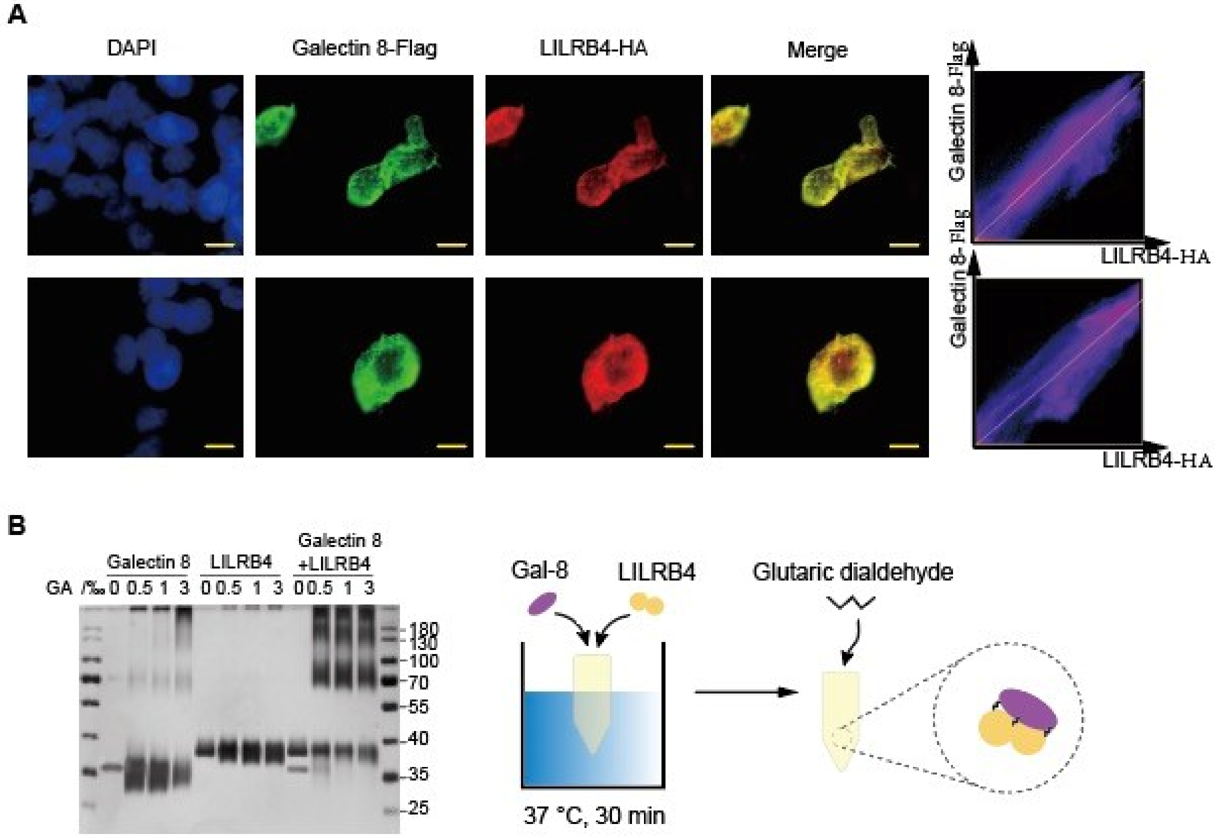


**Supplementary Figure 2.** (A) Colocalization of Galectin 8 and LILRB4. (B) PAGE of Galectin 8 and LILRB4 crosslinked with Glutaric dialdehyde shows direct interaction.

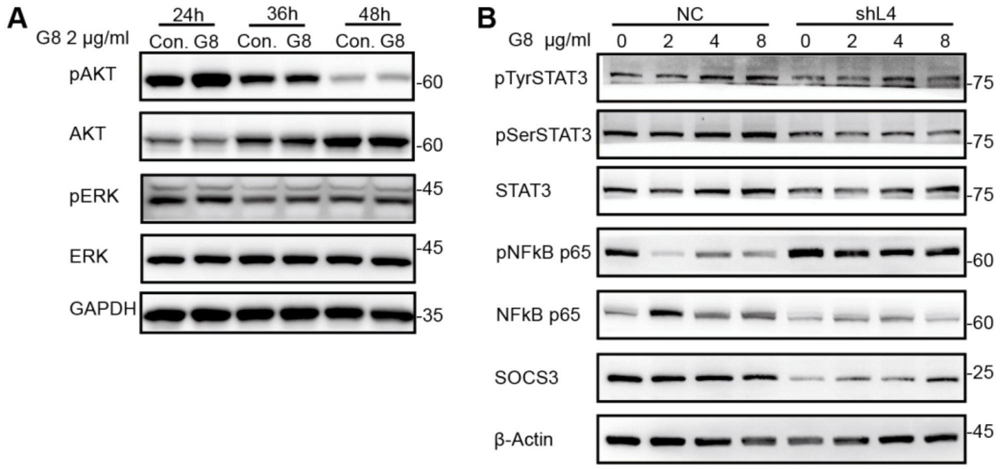


**Supplementary Figure 3. (**A) Potential downstream signals of PTPs including Erk and Akt remained unaffected in THP-1 cells. (B) Concentration scale of Galectin 8 affects major factors in THP-1 cells.

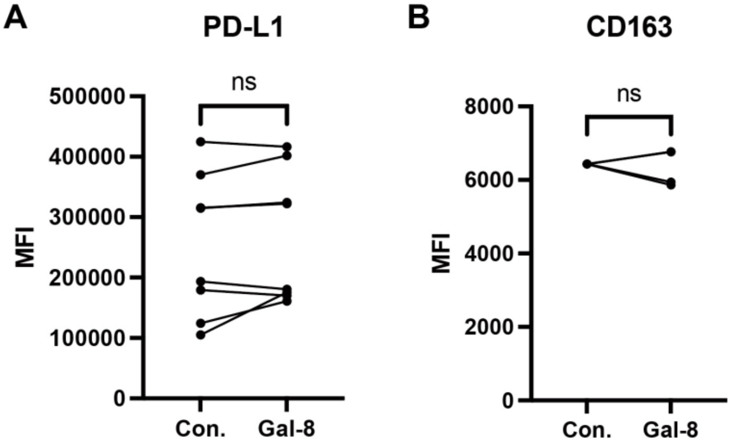


**Supplementary Figure 4.** **(**A, B) Potential ADAM17 targeting membrane proteins including PD-L1 and CD163 was not significantly modulated.

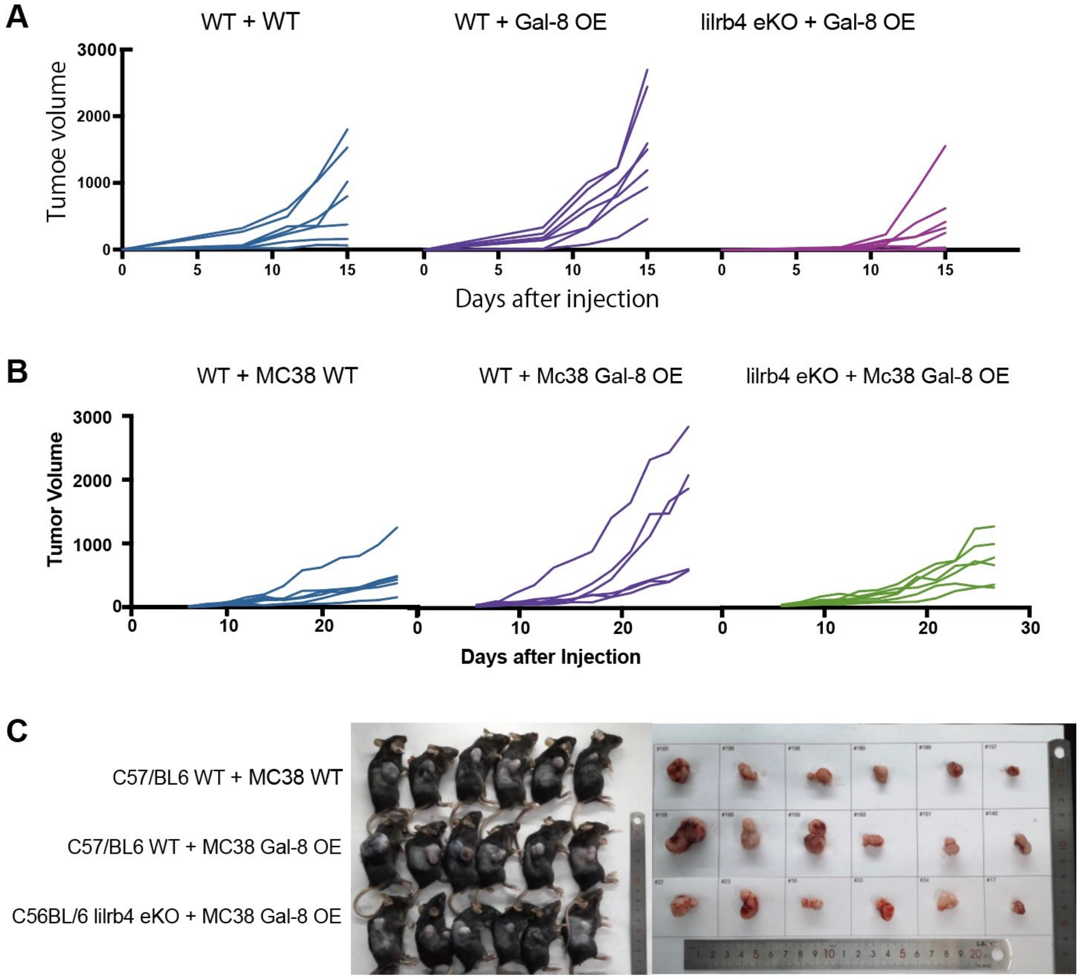


**Supplementary Figure 5.** (A) Tumor volume of each mouse transplanted with B16 tumor. (B) Tumor volume of each mouse transplanted with B16 tumor. (C) Photo of Mc38 tumor in vivo and ex vivo on day 28.

### Supplementary Table 1. Significantly upregulated genes

| Gene Name | Description | Reference |
|---|---|---|
| **CXCL1/2** | CXCL1 and CXCL2 promoted mo-MDSC generation by favoring the differentiation of bone marrow cells in tumor-bearing conditions. | [1, 2] |
| **CCL7** | The level of CCL7 in blood was positively related to the number of Mo-MDSCs in CCR patients, and highly linked with the short-time recurrence and distant metastasis. | [3] |
| **FCN1, VCAN** | As members of S100A family, FCN1 and VCAN were upregulated in MDSC-like cells and acknowledged as an MDSC marker. | [4] |
| **C3** | The C3(-/-) HpSCs lost their ability to induce MDSCs. | [5] |
| **C300e** | The ligation of CD300e in monocytes hampers the expression of HLA II, negatively impacting their capacity to activate T cells in an antigen-specific manner. | [6] |
| **MMP14** | MMP14 was a critical molecule in MDSC migration by CXCL10/TLR4 signaling. | [7] |
| **FPR1** | FPR1 was upregulated upon MDSC co-culturing of tumor bearing mice. | [8] |
| **VSIG4** | VSIG4 is a B7 family-related protein expressed on APCs that strongly inhibits murine and human T cell proliferation and IL-2 production. | [9] |
| **SEMA6B** | SEMA6B expression was positively correlated with infiltrating levels MDSCs and with diverse marker sets of immunosuppressive cells in CRC, predicting poor prognosis. | [10] |
| **THBS1** | THBS1 triggers macrophage IL-10 production. | [11] |
| **CXCL5** | CXCL5 is associated with MDSC accumulation and predicts poor prognosis. | [12, 13] |
| **CCL2** | CCL2 promotes MDSC population and function. | [14] |
| **S100A8/9/12** | Members of the S100 protein family-S100A8, S100A9 and S100A12 - specifically expressed in CD14(+) HLA-DR(-/low) MDSC. | [15] |
| **PPBP** | Monocytes secret PPBP(CXCL7) to promote cancer progression. | [16] |
| **IL-6** | The cytokine interleukin (IL)-6 was found to be a crucial regulator of MDSC accumulation and activation as well as a factor, stimulating tumor cell proliferation, survival, invasiveness and metastasis. | [17] |
| **OSM** | Myeloid-derived OSM reprogrammed fibroblasts to a more contractile and tumorigenic phenotype and elicited the secretion of VEGF and proinflammatory chemokines CXCL1 and CXCL16, leading to increased myeloid cell recruitment. | [18] |
| **OLFML2B** | OLFML2B was positively correlated with the degree of macrophage infiltration and highly co-expressed with tumor-associated macrophage markers. | [19] |
| **CHI3L1** | CHI3L1 contributes to tumor progression by stimulating the PD-1/PD-L1 axis and other checkpoint molecules. | [20] |
| **CA12** | The accumulation of CA12+ macrophages in tumor tissues was associated with increased tumor metastatic potential and reduced survival. | [21] |
| **CEBPB** | C/EBPβ induces transcription factor NFI-A to support the generation and expansion of MDSCs in the bone marrow and spleens of septic mice. | [22] |
| **IRAK3** | IRAK-M (IRAK3) negatively regulates monocytes and macrophage activation. | [23] |
| **IL1A** | IL-1 pathway blockade improves response to therapy through elimination of MDSCs. | [24] |
| **CD163** | The presence of CD163 on classical monocytes suggests that these cells have anti-inflammatory properties. | [25] |
| **LAIR1** | High level of LAIR1 was detected on TAMs and monocytic MDSCs. In vitro, blocking LAIR1 enhanced T-cell activation and proliferation, decreased the expression of M2 markers and increased the level of co-stimulatory proteins. | [26] |

| | | |
|---|---|---|
| **CXCL3/6** | CXCL3-CXCR2 axis promotes immune suppression and metastasis through MDSC. CXCL6 also binds CXCR2. | [27, 28] |
| **MERTK** | MDSCs dramatically upregulated MERTK and its ligand in tumor-bearing mice, leading to immune suppression and resistance to anti-PD-1 therapy. | [29] |
| **AQP9** | Regulatory T cells (Tregs), macrophages M2, macrophages M0, CD4+ T cells, and neutrophils were positively correlated with *AQP9* expression. | [30] |
| **VNN2** | VNN2 was identified as an MDSC biomarker on peripheral blood $CD14^{+}$ monocytes. | [31] |
| **LILRB1/2** | LILRB1 and LILRB2 both interact with HLA-G and cause T cell inhibition through MDSC functioning. | [32] |
| **MMP8** | Persistence of MDSC also may be associated with increased expression of proangiogenic proteins, such as MMP8 produced by tumor stromal cells or infiltrating MDSCs. | [33] |
| **IL10** | IL-10 promotes MDSC development and production of IL-10 by MDSC induces Treg. | [34, 35] |

### Table 2. Significantly downregulated genes

| Gene Name | Description | Reference |
|---|---|---|
| **CD1A/B/C/E** | CD1a, CD1b and CD1c capture distinct classes of self- and mycobacterial antigens. CD1+ dendritic cells are involved in the first steps of the primary immune response in a number of malignancies and the lack of CD1a+ cells corelates with malignant evolution. CD1e participates in lipid antigen pre-sentation without interacting with the T-cell receptor. | [36-38] |
| **CCL17** | CCL17 was reported to promote inflammatory reaction and suppress Treg recruitment. | [39, 40] |
| **MMP12** | MDSCs accumulated stronger in the bone marrow of $MMP12^{-/-}$ mice compared with wild type $MMP12^{+/+}$ mice. | [41] |
| **IFIT1** | Ifit1 was expressed on macrophages of mice and silencing it suppressed LPS induced activation. | [42] |
| **RSAD2** | Mature DCs lost their anti-tumor efficacy upon Rsad2 knockdown. | [43] |
| **IFI44, IFI44L** | Low expression of IFI44L levels also correlated with larger tumor size, disease relapse, advanced stages, and poor clinical survival in HCC patients. | [44] |
| **ADAM19** | ADAM19 is a marker of human DCs and correlates with tumor immune cell infiltration. | [45, 46] |
| **HLA-DRA** | MDSC was defined as CD11b+CD33+HLA-DR-/lo cells. Generally, MHC II was downregulated in MDSC compared with macrophages. | [47] |
| **USP18** | Usp18 promotes conventional CD11b+ DC development through inhibiting Type I interferon signaling. | [48] |
| **CLDN1** | CLDN10 correlates with lymph node metastasis but predicts favorable prognosis. | [49] |
| **DNASE1L3** | Down-regulation of DNASE1L3 may participate in immune escape via CCR7/CCL19 axis. | [50] |
| **CLEC10A** | CLEC10A enhances cytokine secretion of DCs and correlates with tumor immune infiltrates. | [51, 52] |
| **LIPA** | Loss of the lysosomal acid lipase function leads to expansion of myeloid-derived suppressive cells (MDSCs) that cause myeloproliferative neoplasm. | [53] |
| **STAT1** | In STAT1-/- mice, increase in the frequency of the MDSC-like cells and lung IL-10 levels was observed upon infection. | [54] |
| **IRF7** | IRF7 deficiency caused significant elevation of G-MDSCs, and therefore enhanced tumor growth and metastasis in mice. | [55] |

| | | |
|---|---|---|
| **ECM1** | The macrophage-specific knockout of ECM1 resulted in increased arginase 1 (ARG1) expression and impaired polarization into the M1 macrophage phenotype after lipopolysaccharide (LPS) treatment. | [56] |
